# Disruption of Epithelium Integrity by Inflammation-Associated Fibroblasts through Prostaglandin Signaling

**DOI:** 10.1101/2023.09.28.560060

**Authors:** Yi Dong, Blake A. Johnson, Linhao Ruan, Maged Zeineldin, Albert Z. Liu, Sumana Raychaudhuri, Ian Chiu, Jin Zhu, Barbara Smith, Nan Zhao, Peter Searson, Shigeki Watanabe, Mark Donowitz, Tatianna C. Larman, Rong Li

**Affiliations:** Department of Cell Biology, Johns Hopkins School of Medicine; Baltimore, MD, 21205, U.S.A; Department of Pathology, Division of GI/Liver Pathology, Johns Hopkins School of Medicine; Baltimore, MD, 21205, U.S.A; Mechanobiology Institute and Department of Biological Sciences, National University of Singapore; Singapore; Microscope Facility, Johns Hopkins School of Medicine; Baltimore, MD, 21205, U.S.A; Institute for Nanobiotechnology, Johns Hopkins University; Baltimore, Maryland, 21218, U.S.A; Department of Materials Science and Engineering, Johns Hopkins University; Baltimore, MD, 21218, U.S.A; Department of Medicine, Division of Gastroenterology, Johns Hopkins School of Medicine; Baltimore, MD, 21205, U.S.A; Department of Physiology, Johns Hopkins School of Medicine; Baltimore, MD, 21205, U.S.A; Department of Biological Sciences, National University of Singapore; Singapore

## Abstract

Inflammation-associated fibroblasts (IAFs) are associated with the progression and drug resistance of chronic inflammatory diseases such as inflammatory bowel disease (IBD), but their direct impact on epithelial function and architecture is unknown. In this study, we developed an *in vitro* model whereby human colon fibroblasts are induced to become IAFs by specific cytokines and recapitulate key features of IAFs *in vivo*. When co-cultured with patient-derived colon organoids (colonoids), IAFs induced rapid colonoid swelling and barrier disruption due to swelling and rupture of individual epithelial cells. Epithelial cells co-cultured with IAFs also exhibit increased DNA damage, mitotic errors, and proliferation arrest. These IAF-induced epithelial defects are mediated through a paracrine pathway involving prostaglandin E2 (PGE2) and the PGE2 receptor EP4, leading to PKA-dependent activation of the CFTR chloride channel. Importantly, EP4-specific chemical inhibitors effectively prevented colonoid swelling and restored normal proliferation and genome stability of IAF-exposed epithelial cells. These findings reveal a mechanism by which IAFs could promote and perpetuate IBD and suggest a potential treatment to mitigate inflammation-associated epithelial injury.

**Teaser:** Inflammation-associated fibroblasts compromise colon epithelial barrier integrity and genome stability via PGE2-EP4 signaling.

## Introduction

Fibroblasts are a heterogeneous group of stromal cells that contribute to tissue architecture and support homeostasis of resident cells (*1*). Recent advances in single-cell multi-omics have provided insights into the diversity and functions of fibroblasts in normal and disease-affected tissues (*2, 3*). Fibroblasts help maintain tissue integrity and homeostasis by secreting inflammatory mediators, producing growth factors and extracellular matrix (ECM) components, and facilitating the remodeling of tissue architecture after injury (*4*). However, under pathological conditions such as chronic inflammation, fibroblasts can be dysregulated to become inflammation-associated fibroblasts (IAFs) and contribute to pathogenic tissue fibrosis and scarring (*2, 5*) and potentially other short- and long-term consequences.

Inflammatory bowel disease (IBD) is a chronic inflammatory condition affecting the intestine. Depending on the constellation of clinical symptoms and pattern of injury in the tubular gastrointestinal tract, IBD is partitioned into Crohn’s disease (CD) and ulcerative colitis (UC). Its etiology is complex and multifactorial, but a key contributor to IBD pathogenesis is chronic epithelial barrier dysfunction that can instigate and propagate excessive immune responses (*6*). Chronic mucosal injury and repair can lead to mucosal remodeling including crypt architectural distortion, fibrosis, expanded lamina propria chronic inflammation, and epithelial metaplasia (*7*). IBD patients suffer from chronic diarrhea, fibrostenotic disease, and fissures, and carry increased risk of colitis-associated dysplasia and colorectal cancer (CAC) (*8-10*). CAC is characterized by early *TP53* mutations and widespread chromosome instability (CIN) (*11, 12*). DNA damage and aneuploidy are observed even in non-dysplastic IBD epithelium and likely plays a key role in CAC evolution (*13*). A recent human study reported that IBD colonic epithelium accrued twice the number of gene mutations and aneuploidy than normal colon epithelium (*10*). However, mechanisms by which the chronic inflammatory microenvironment in IBD promotes genome instability are poorly understood.

Because fibroblasts regulate the stemness, wound healing, and differentiation of intestinal epithelial cells (*14*), IAFs may play a role in IBD epithelial dysfunction. Recent scRNAseq studies have defined IL13RA2^+^ IL11^+^ fibroblasts as IAFs in IBD (*15*). These IAFs showed a strong association with immune signaling, extracellular matrix (ECM) remodeling, and epithelial regulation (*16*). Importantly, these IAFs were linked to resistance to TNF blocking agents commonly used to treat IBD (*15, 17-19*). Recent clinical studies have shown that reduction of IAFs is strongly associated with the responsiveness of IBD biologics, which mostly target immune cells (*17, 20, 21*). Moreover, animal studies have shown that IAFs correlate with poor prognosis of CAC (*22, 23*). However, how IAFs interact with colon epithelia remains elusive. In this study, we develop an *in vitro* model of human colon derived IAFs and use it to define cellular and molecular interactions between colon IAFs and epithelial cells. Our experimental findings uncover a paracrine pathway by which IAFs promote trans-epithelial fluid secretion, leading to impaired barrier function and cellular and genomic abnormalities in colon epithelium.

## Results

### *In vitro* induction of IAFs from patient-derived fibroblasts

We first attempted to obtain IAFs from surgically resected colon tissues of IBD patients and from normal controls (Fig. 1A, S1A). We confirmed the presence of IL13RA2^+^ Vim^+^ IAFs, which were enriched in surgical resections from IBD patients (*15*) (Fig. 1B, S1B). We then used a passive selection approach to enrich bulk fibroblasts (see details in Methods) from normal, UC, CD samples (Supplemental Table 1). However, IL13RA2 immunoblotting of protein lysates from IBD-derived early passage bulk fibroblasts was negative, suggesting that IAFs were no longer present (Fig. 1C, S1C-F). To further investigate this, we cultured IL13RA2^+^ fibroblasts directly sorted from fresh IBD patient tissue, but once again IL13RA2 expression was lost after a short expansion (4 passages) *in vitro* (Fig. 1C). We hypothesized that IL13RA2^+^ fibroblasts require a continuous pro-inflammatory environment to maintain their identity. We therefore treated bulk fibroblasts with IL1β, IL4, and TNFα, common cytokines detected in IBD colon mucosa (*24-26*). Indeed, cultured fibroblasts regained IL13RA2 expression after cytokine treatment (Fig. 1C, S1C-F), similar to a previous report in nasal polyp fibroblasts (*27*). This suggests that IL13RA2^+^ fibroblasts may require continuous cytokine stimulation to maintain IAF characteristics.

**Figure 1.**
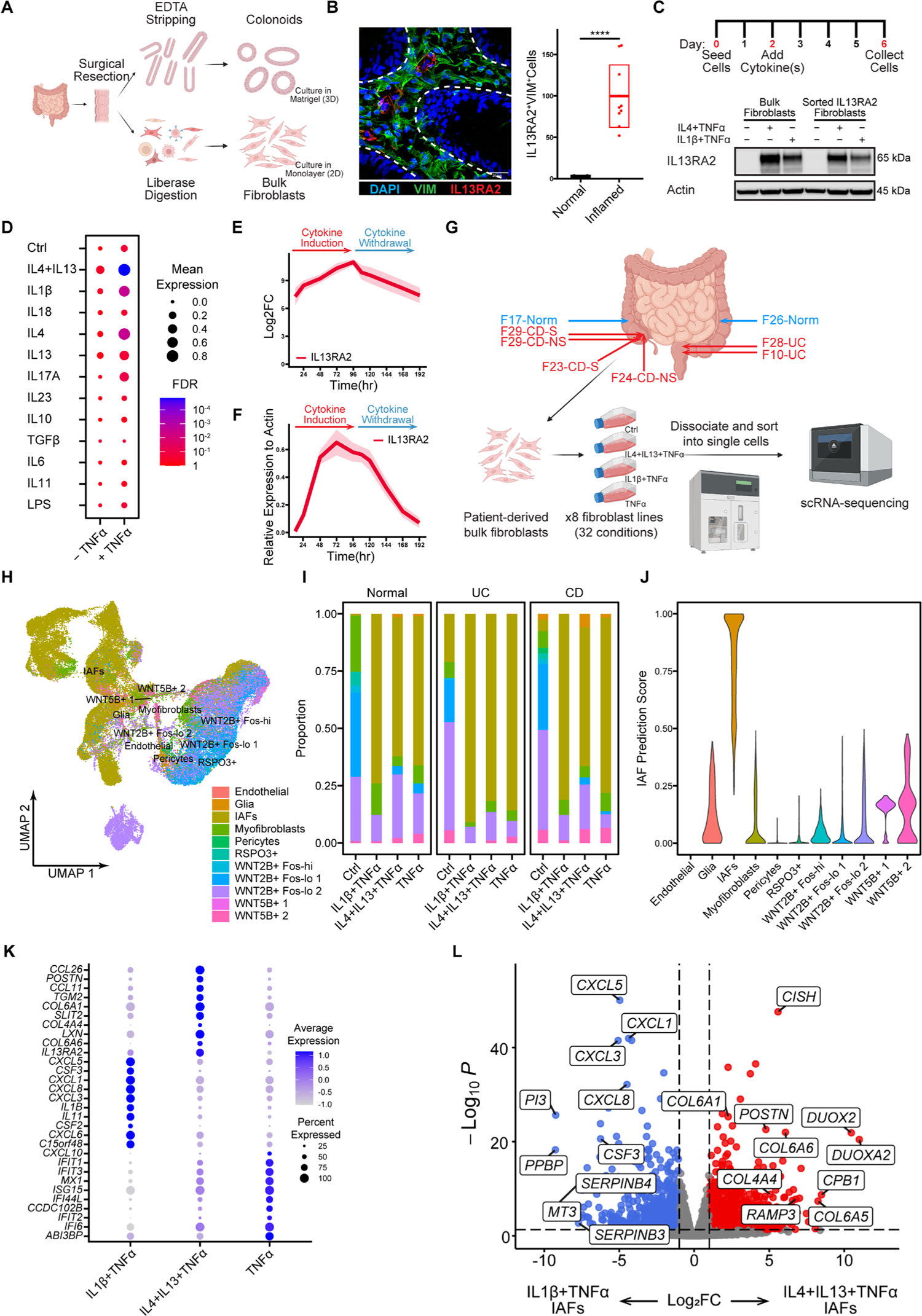
Generation of patient-derived fibroblasts and induction of IAFs. A. Schematic representation depicting the workflow for generating patient-derived bulk fibroblasts and colonoids. Detailed sample information is listed in supplemental table 1 and details are described in Methods. B. Visualization of IAFs in human colonic mucosa. Left: Immunofluorescent image of a frozen human tissue section illustrating the presence of VIM^+^IL13RA2^+^ IAFs; Scale bar = 20 µm. Right: Number of IAFs in the field of view. n = 10 for IBD sections (7 UC, 3 CD) and n = 7 for normal sections. ****p < 0.0001, t-test, error box: STD. C. *In vitro* expanded fibroblasts do not express IL13RA2 without cytokine stimulation. Top: Schematic representation depicting the workflow for activating fibroblasts using cytokine cocktails. Blot: Immunoblot showing the expression of IL13RA2 in *in vitro* expanded fibroblasts treated with or without different cytokines. D. Summary of IL13RA2 expression in fibroblasts treated with different cytokine cocktails. Data from immunoblots of 11 patient derived fibroblast lines (4 UC, 4 CD,3 normal). Mean expression (dot size) and False Discovery Rate (FDR, dot color) of IL13RA2 protein expression are listed in the presence of different cytokines (rows) with or without the addition of TNFα (columns). E. mRNA expression of IAF marker *IL13RA2* is inducible with IL4+IL13+TNFα cocktail. Time course of qPCR assay showing the mRNA expression levels of *Il13RA2* during cytokine induction and withdrawal. Data are summarized from 5 patient derived fibroblast lines (2 UC, 2CD, 1 normal). The line represents the mean level of mRNA expression, and the shaded area represents SEM. F. Protein expression of IAF marker IL13RA2 is inducible with IL4+IL13+TNFα cocktail. Time course of immunoblot assay showing the protein expression levels of IL13RA2 during cytokine induction and cytokine withdrawal. Fibroblasts were first treated with IL4+IL13+TNFα for 96hrs. After washing, cells were cultured in fresh media without cytokines for 96hrs. Data are summarized from 5 patient derived fibroblast lines (2 UC, 2CD, 1 normal). The line represents the relative mean expression level of IL13RA2 compared to β-actin, and the shaded area represents SEM. G. Schematic representation of the workflow for collecting, expanding, and treating patient-derived fibroblasts with different cytokine cocktails for scRNAseq. Details are described in Methods. H. Cell census and cross reference of patient derived fibroblasts. Shown is a uniform manifold approximation and projection (UMAP) of cells labelled by cell subsets. Cells were annotated using a cross reference approach from a published dataset (*15, 29*). I. IAFs are enriched after cytokine activation. Bar plots showing the proportion of different fibroblast subsets in control, IL1β+TNFα, IL4+IL13+TNFα, and TNFα fibroblasts. J. IAF census and similarity score. Violin plot showing the predicted IAF score of fibroblasts in each subset. Details are described in Methods. A similarity prediction score is assigned ranging between 0 and 1. K. Subset-specific markers of different cytokine treated fibroblasts. Shown are the differentially expressed genes (DEGs) of fibroblasts treated with different fibroblasts (IL1β+TNFα, IL4+IL13+TNFα, and TNFα). The percentage of cells expressing (dot size) and the mean expression level (dot color) of selected genes (rows) across subsets (columns) are shown. L. DEGs that distinguish IL1β+TNFα vs. IL4+IL13+TNFα IAFs. Volcano plot shows the DEGs between IL1β+TNFα (left) and IL4+IL13+TNFα (right) activated IAFs. 7,529 and 6,465 cells were analyzed from each subset and 27445 variables were identified.

To test the above hypothesis, we screened a panel of IBD-associated cytokines and inflammatory factors to determine which led to robust upregulation of IL13RA2 in 4 UC, 4 CD and 3 normal fibroblast lines (*24-26*). Most cytokines were insufficient to induce IL13RA2 upregulation individually, although IL1β, IL4, IL13, and TNFα led to moderate upregulation (Fig 1D, S1G). When combining Th1 cytokines IL1β and TNFα, two classic cytokines that activate fibroblasts in cancer (*28*), IL13RA2 expression was markedly upregulated (Fig. 1D). However, Th2 cytokines IL-4 and IL-13, in combination with TNFα, induced the strongest IL13RA2 expression (Fig 1D). To determine the kinetics of IL13RA2 induction and turnover, we used the IL4+IL13+TNFα cytokine cocktail and performed a time course experiment with 4 days of cytokine treatment followed by 4 days of cytokine withdrawal. Both the mRNA and protein level of IL13RA2 increased with cytokine treatment and decreased upon cytokine withdrawal (Fig 1E, S1F). These data show that IL13RA2+ fibroblasts can be induced from both normal and IBD colon-derived bulk fibroblasts, but their maintenance requires continuous presence of the cytokines.

To further evaluate the similarity between cytokine-activated fibroblasts as described above and IAFs *in vivo*, we performed single-cell RNA sequencing (scRNAseq) of patient-derived bulk fibroblasts obtained from 2 UC samples, 2 CD non-stenotic samples, 2 CD stenotic samples, and 2 healthy controls (Fig. 1G, S1H). The fibroblasts were treated for 4 days with either TNFα alone, IL4+IL13+TNFα, or IL1β+TNFα. Including control fibroblasts, 60,845 cells were recovered with mean reads of 75,614. Using a published reference mapping method (*29*), we compared single-cell gene expression profiles with a published UC atlas database (*15*) and confirmed that our cytokine-activated fibroblasts were enriched to the IAF category (Fig. 1H, S1I, S1J), compared to other populations (Fig. 1J, S1K). Also consistent with the lack of IL13RA2 protein expression in early-passage non-cytokine-treated bulk fibroblasts (Fig. 1C), cells mapping to IAF were strongly enriched after cytokine induction (Fig. 1I). Interestingly, fibroblasts treated with different cytokine cocktails showed unique gene expression signatures (Fig. 1K, S1L). Comparing differentially expressed genes (DEGs) in IL1β+TNFα activated fibroblasts revealed enrichment of genes involved in chemotaxis (e.g., *CXCL1*, *CXCL3*, *CXCL6*, *CXCL8*, *CXCL10*, *CSF2*, *CSF3*) (Fig. 1K, 1L). By contrast, IL4+IL13+TNFα-activated fibroblasts showed upregulation of ECM genes (e.g., *COL4A4*, *COL6A1*, *COL6A6*, *POSTN*), whereas fibroblasts treated with TNFα alone showed enrichment for genes downstream of interferon signaling (e.g., *IFIT1*, *2*, *3*, *6*) (Fig. 1K). Although gene set enrichment analysis (GSEA) suggested that fibroblasts treated with either IL1β+TNFα or IL4+IL13+TNFα showed enrichment in pathways associated with immune responses (Fig. S1M, S1N), IL4+IL13+TNFα-activated fibroblasts showed significant enrichment in genes associated with collagen processing and ECM remodeling (Fig. 1L). These signatures are also significantly enriched in GSEA analysis of IAFs from IBD patients (Fig. S1O). Additionally, IL4+IL13+TNFα-induced IAFs showed significantly higher expression of many previously-reported IBD-associated IAF transcriptional signatures (*15, 17, 30-33*) compared to compared to IL1β+TNFα-treated IAFs (Fig. S1P). The IL4+IL13+TNFα-activated fibroblasts were therefore used as IAFs for the subsequent experiments in this study.

### IAFs induce colonoid swelling via paracrine signaling

Fibroblasts are known to regulate the morphology, expansion, and differentiation of epithelial cells in the intestinal crypt (*18*). To investigate how IAFs affect colon epithelial organization and growth (Fig. S2A), we established a co-culture model combining either IAFs or normal fibroblasts (NFs) with human colonoids (Fig. 1A, 2A, see Methods for details). To generate human colonoids, the EDTA based stripping method was used to enrich crypts. Dissociated crypts were seeded in Matrigel and cultured for 4 days to form colonoids. In the meantime, IAFs were induced using IL4+IL13+TNFα for 4 days. Normal fibroblasts (NFs) or IAFs were then co-cultured with colonoids outside of the Matrigel. Importantly, IAF induction was followed by cytokine washout prior to co-culturing with normal colonoids to avoid any direct effect of the cytokines on colonoids, but the duration of the co-culture was only 24 hrs before IAFs lost their characteristics as shown in Figure 1F. By using live-cell imaging, we observed that the luminal volume of colonoids increased dramatically when co-cultured with IAFs but not with NFs (Fig. 2B, 2C, supplement movie 1). The same phenomenon was observed when transwells were used as a physical barrier between IAFs and colonoids (Fig. S2B, S2C), suggesting that colonoid swelling did not require direct contact between IAFs and epithelial cells.

**Figure 2.**
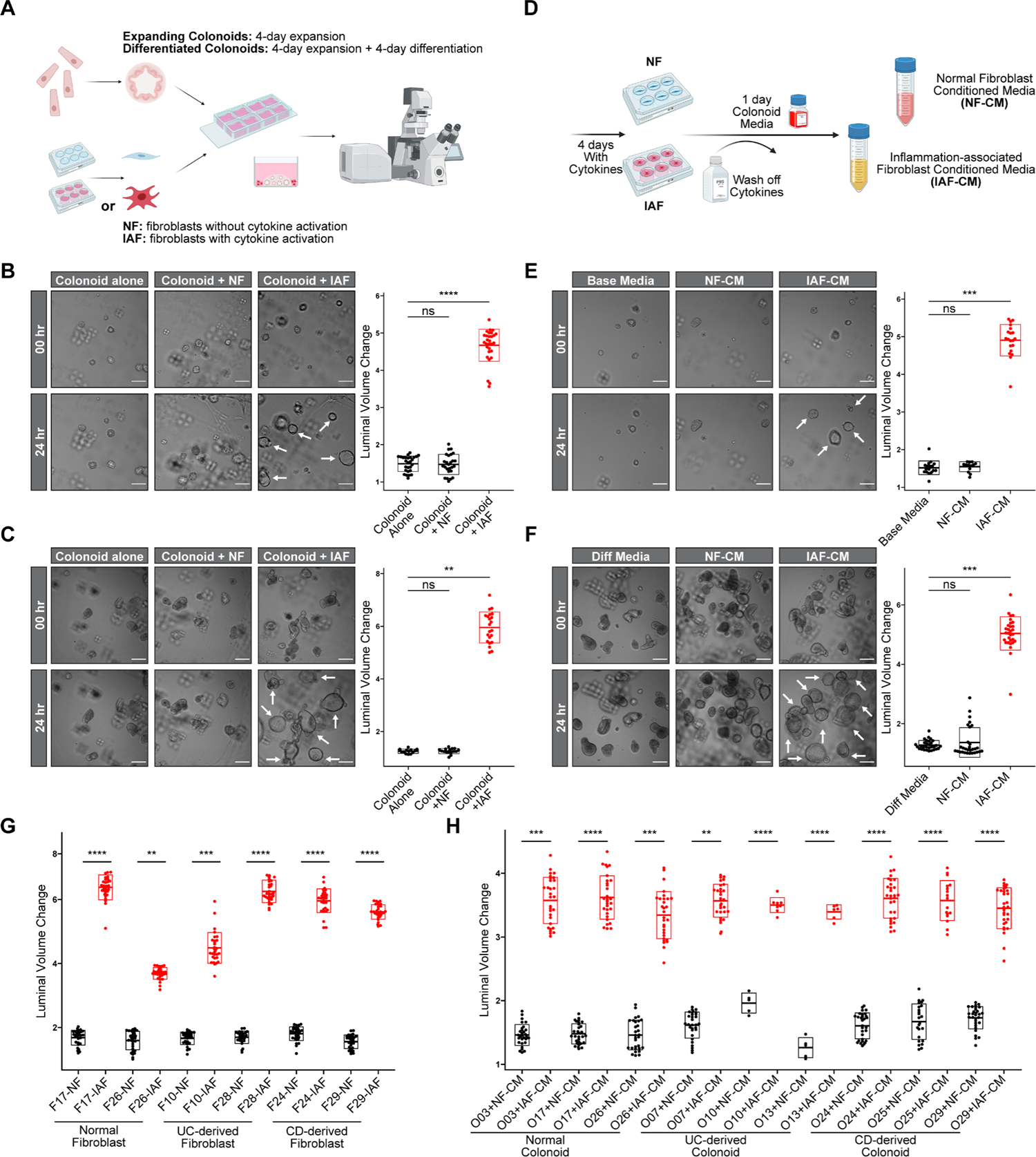
IAFs induce colonoid swelling through paracrine signaling. A. Schematic representation of the workflow for colonoid-fibroblast co-culture experiments. Details are described in Methods. B. IAFs induce colonoid swelling in expanding colonoids. Left: Representative images of expanding colonoids co-cultured with normal (NF) or cytokine activated (IAF) fibroblasts. Arrows point to swelling colonoids co-cultured with IAFs. Images were taken at the start of and 24 hrs post co-culture. Scale bars, 100 µm. Right: Box plots showing the level of luminal volume changes of expanding colonoids co-cultured with NF or IAF. Three independent experiments were performed with consistent results. Data shown is from a single independent experiment, with 27, 26, and 29 colonoids measured per condition (from left to right). Each dot represents an individual colonoid; ****p < 0.0001, ns not significant, t-test, bar box represents mean ± STD. C. IAFs induce colonoid swelling in differentiated colonoids. Left: Representative images of differentiated colonoids co-cultured with NFs or IAFs. Arrows point to swelling colonoids co-cultured with IAFs. Images were taken at the start of and 24 hrs post co-culture. Scale bars, 200 µm. Right: Right: Box plots showing the level of luminal volume changes of differentiated colonoids co-cultured with NF or IAF. Three independent experiments were performed with consistent results. Data shown is from a single independent experiment, with 13, 16, and 22 colonoids measured per condition (from left to right). Each dot represents an individual colonoid; **p < 0.01, ns not significant, t-test, bar box represents mean ± STD. D. Schematic representation of the workflow for generating fibroblast conditioned media (CM) for colonoid-fibroblast co-culture experiments. E. IAF-CM induces colonoid swelling in expanding colonoids. Left: Representative images of expanding colonoids cultured in normal (NF-CM) or cytokine-activated (IAF-CM) fibroblast conditioned media. Arrows point to swelling colonoids under IAF-CM. Images were taken at the beginning of and 24 hrs post colonoid culture in CM. Scale bars, 100 µm. Right: Box plots showing the level of luminal volume changes of expanding colonoids cultured using NF-CM or IAF-CM. Three independent experiments were performed with consistent results. Data shown is from a single independent experiment, with 13, 17, and 20 colonoids measured per condition (from left to right). Each dot represents an individual colonoid; ***p < 0.001, ns not significant, t-test, bar box represents mean ± STD. F. IAF-CM induces colonoid swelling in differentiated colonoids. Left: Representative images of differentiated colonoids cultured in normal (NF-CM) or cytokine-activated (IAF-CM) fibroblast conditioned media. Arrows point to swelling colonoids under IAF-CM. Images are taken at the beginning of and 24 hrs post colonoid culture in CM. Scale bars, 200 µm. Right: Box plots showing the level of luminal volume changes of differentiated colonoids cultured using NF-CM or IAF-CM. Three independent experiments were performed with consistent results. Data shown is from a single independent experiment, with 30 colonoids measured per condition. Each dot represents an individual colonoid; ***p < 0.001, ns not significant, t-test, bar box represents mean ± STD. G. IAFs from multiple fibroblast lines induce colonoid swelling. Box plots showing the level of luminal volume changes of colonoid O03 treated with NF or IAF induced by six different fibroblast lines (2 normal, 2 UC, and 2 CD). Three independent experiments were performed with consistent results. Data shown is from a single independent experiment, with 30 colonoids measured per condition except F17CN-NF (29), F10UC-NF (28), F28UC-NF (26). Each dot represents an individual colonoid; ****p < 0.0001, ***p < 0.001, **p < 0.01, t-test, bar box represents mean ± STD. H. Multiple colonoid lines respond to IAF-CM induced swelling. Box plots showing the level of luminal volume changes of nine colonoid lines (3 normal, 3UC, and 3CD) treated with NF-CM or IAF-CM derived from fibroblast F26. Three independent experiments were performed with consistent results. Data shown is from a single independent experiment, with 30 colonoids measured per condition except O10UC+NF-CM and IAF-CM (5 and 8), O13UC+NF-CM and IAF-CM (5, and 6), and O25CD+NF-CM and IAF-CM (24, and 17). Each dot represents an individual colonoid; ****p < 0.0001, ***p < 0.001, **p < 0.01, t-test, bar box represents mean ± STD.

To directly test whether IAF-secreted paracrine factors caused colonoid swelling, we added colonoid culture media to fibroblasts and collected conditioned media (CM) from 6 different fibroblast lines (2 UC, 2 CD, 2 Normal), with or without cytokine induction (Fig. 2D, see Methods for details). We first ruled out the possibility that colonoid swelling was induced by traceable amounts of IAF-inducing cytokines (IL1β, IL4, IL13, or TNFα) after cytokine washout by treating colonoids with these cytokines directly (Fig. S2D). CM were then used to culture colonoids (Fig. 2E, 2F). In all colon fibroblast lines tested, CM produced by IAFs (IAF-CM), but not normal fibroblasts (NF-CM), induced colonoid swelling (Fig. 2G). Similar results were obtained when using IAF-CM from one of the fibroblast lines to culture colonoid lines derived from normal, UC, and CD patients (Fig 2H). These results confirmed that IAFs induced colonoid swelling via paracrine signaling, and that cytokine-activation, rather than the source of fibroblasts or colonoids, was critical to induce colonoid swelling. Intriguingly, this aligns with immunofluorescent staining of IBD colonic mucosa which shows that IAFs were scattered in the lamina propria and not directly interacting with colon crypts (Fig. S2A). Because the various fibroblast and colonoid lines behaved similarly in co-culture experiments, we used a representative normal colonoid line (O03) and a representative normal fibroblast line (F26) in subsequent experiments.

### IAF-induced colonoid swelling is associated with barrier leakage and cell rupture

To understand how IAFs induce colonoid swelling, we first considered the possibility of increased epithelial cell proliferation. However, terminally differentiated colonoids similarly swelled in the presence of IAF-CM (Fig. 2F, supplemental movie 2), arguing against this possibility (*34*). Next, we generated colonoids expressing H2B-mNeonGreen and tracked cell proliferation for 24 hours by live imaging. Intriguingly, we observed fewer cell divisions in colonoids cultured in IAF-CM compared to those in NF-CM (Fig. 3A, S3A). The EdU incorporation assay further confirmed reduced proliferation in IAF-CM cultured colonoids compared to those cultured in NF-CM (Fig. 3B, S3B). These results suggest that the IAF-induced colonoid swelling was not due to increased cell proliferation but was instead associated with impaired epithelial growth.

**Figure 3.**
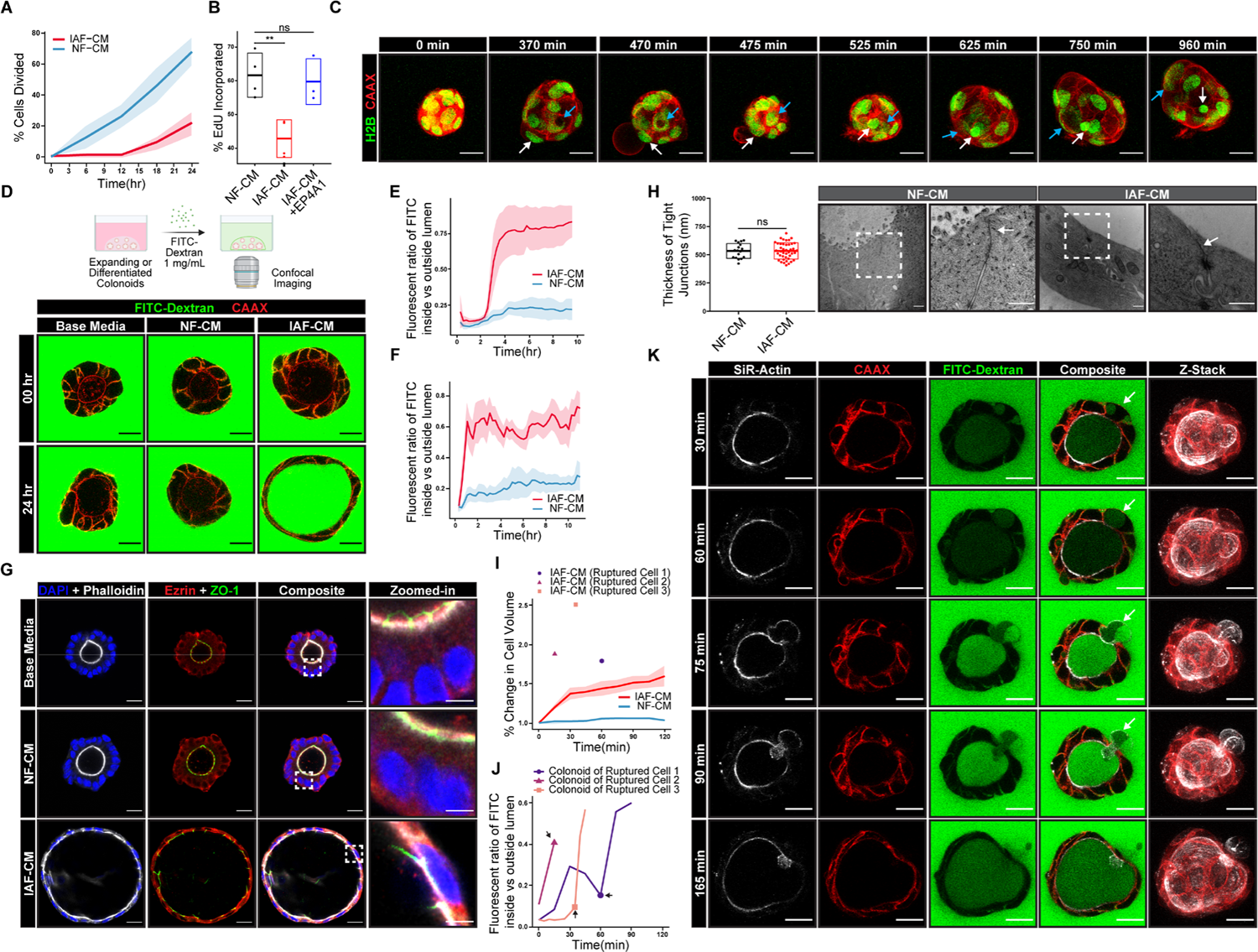
Barrier disruption and cell rupture are observed during IAF-induced colonoid swelling. A. IAFs suppress colonoid proliferation. Summarized data representing the percentage of colonoid cells which divided during 24 hrs in live cell imaging experiments. 6 colonoids (94 cells) cultured with NF-CM and 7 colonoids with IAF-CM (107 cells) were tracked. Solid line represents mean and shaded area represents SEM. B. IAFs inhibit EdU incorporation. Box plot showing the percentage of EdU incorporated in colonoid cells treated with NF-CM, IAF-CM, or IAF-CM with EP4 inhibitor (EP4A1) in the flow cytometry assay. 4, 4, and 3 experiments were performed for each condition. **p < 0.01, ns not significant, t-test, bar box represents mean ± STD. C. IAF-CM induces both colonoid swelling and cell swelling. Representative images showing the change of a colonoid treated with IAF-CM. White arrows point to an individual cell that underwent swelling and rupture. Its nucleus was ejected into the lumen. Blue arrows point to another cell that had a large vacuole in the cell. The vacuole displaced the nucleus and deformed its shape. D. IAF-CM increases the permeability of colonoids. Left: Schematic representation of the workflow used to examine colonoid barrier function. Right: Representative images showing that IAF-CM but not NF-CM increased the permeability of colonoids. Scale bar, 20 µm. E. IAF-CM increases the permeability of expanding colonoids. Line plot shows the permeability of colonoids after IAF-CM or NF-CM culture. Live cell imaging was performed for 10 hrs as described in (L). We monitored n = 6 IAF-CM-treated and n = 6 NF-CM-treated colonoids. Solid lines indicate mean and shaded areas represent SEM. F. IAF-CM increases the permeability of differentiated colonoids. Line plot showing the permeability of colonoids after IAF-CM or NF-CM culture. Live cell imaging was performed for 10 hrs as described in (L). We monitored n = 4 for IAF-CM-treated and n = 5 NF-CM-treated colonoids. Solid lines indicate mean and shaded areas SEM. G. IAF-CM cultured colonoids retain tight junctions and polarity. Representative immunofluorescent images of expanding colonoids cultured in NF-CM or IAF-CM fibroblast conditioned media. Zoomed in images of the boxed areas are shown in the right. Scale bars = 20 µm in original images and bars = 5 µm in zoomed-in images. H. IAF-CM did not affect the intercellular thickness of tight junctions. Left: Box plots showing the intercellular thickness of tight junctions. Data were quantified from TEM images. n = 42 junctions quantified from IAF-CM cultured colonoids and n = 17 from NF-CM cultured colonoids. Right: Representative TEM images of colonoid treated with IAF-CM or NF-CM for 6 hrs. White arrows indicate tight junctions. Scale bar, 500 nm. I. IAF-CM increased cell volume and led to cell rupture in some cells. Line plots show the change in volume over time of individual cells in colonoids treated with IAF-CM or NF-CM. Circle, triangle, and rectangle symbols represent volume changes of three IAF-CM cultured cells when they ruptured. N=10 cells for both IAF-CM treated and NF-CM cultured colonoids. Solid lines represent mean value and shaded areas represent SEM. J. IAF-CM-induced cell rupture was associated with barrier disruption. Line plot showing the fluorescent ratio of FITC inside vs outside colonoid lumen in an FITC-Dextran permeability assay. These three colonoids were cultured with IAF-CM as described in Fig 3I. Circle, triangle, and rectangle dots represent the same cells described in Fig 3I and the timepoints when the swelling cells ruptured. K. IAF-CM induced colonoid swelling, rupture, and barrier disruption. Representative live images of a colonoid in IAF-CM culture. Time course of a colonoid cell (white arrow) exhibited increased volume within the cell and eventually ruptures. FITC-dextran leaks into the lumen because of the rupture.

Time-lapse imaging showed that colonoids began to increase in luminal volume within minutes of IAF-CM treatment, followed by a sudden volume reduction a few hours later (Fig. 3C, Supplement movie 3). Whereas the volume increase could be due to transepithelial fluid secretion, the collapse could be related to rupture of the epithelial barrier (Fig. 3C). To test this, we assayed epithelial permeability by adding 4 kDa FITC-dextran into the media (Fig. 3D). In NF-CM, there was no significant leakage of FITC-dextran into the colonoid lumen (*35*). In contrast, both non-differentiated and terminally differentiated colonoids were susceptible to FITC-dextran leakage into the lumen when treated with IAF-CM (Fig. 3E, 3F, S3D, S3E). These observations suggest that IAF paracrine signaling can disrupt the barrier function of colon epithelium.

The barrier function of epithelia requires apical-basal polarity and polarized formation of tight junctions (*36*). Immunofluorescent staining (IF) of both tight junctions (stained using anti-ZO-1) and cell polarity markers (actin, stained with phalloidin) revealed that epithelial cells in colonoids cultured in IAF-CM lost their normal columnar morphology and became stretched circumferentially (Fig. 3G, S3C). However, thin-sectioning electron microscopy showed that epithelial junctions appeared largely intact in IAF-CM treated colonoids (Fig 3H, S3F). Similar results were observed in colonoids expressing mNeonGreen::ZO-1 (Fig. S3G-J). These data suggest that IAF-CM unlikely compromised the integrity of tight junctions.

To observe dynamic epithelial changes during IAF-induced colonoid swelling, we performed live-cell imaging using colonoids expressing plasma membrane-anchored red fluorescent protein (mCherry-CAAX) and labelled with SiR-Actin, a live-actin probe (*37*). In conjunction with the FITC-Dextran permeability assay, we observed that individual epithelial cells were larger in colonoids cultured in IAF-CM than those in NF-CM (Fig. 3I), and FITC-Dextran leakage into the cell was associated with dramatic swelling and rupture of individual cells in colonoids (Fig. 3C, 3J), which was associated with a rapid increase of FITC-Dextran signal in the colonoid lumen (Fig. 3K). These observations suggest that cell rupture is a cause of epithelial barrier disruption during swelling.

### IAF-induced colonoid swelling is PKA and CFTR-dependent

The rapid increase of colonoid luminal volume suggests trans-epithelial fluid secretion into the lumen, a process contributing to diarrhea in IBD (*38*). As it was reported that PKA, PKC, and PKG regulate cellular processes in diarrhea (*39-41*), we treated colonoids with corresponding agonists for each of these kinases (Forskolin for PKA; PMA for PKC; 8-br-cGMP for PKG). Colonoids treated with forskolin (*42*), but not PMA or 8-br-cGMP, showed robust swelling (Fig. 4A). We then treated colonoids with four chemically distinct PKA inhibitors (SQ-22536 and KH 7 target adenylyl cyclase, the upstream of PKA; H-89 and A-674563 target PKA) and observed that all four inhibitors inhibited the IAF-CM-induced colonoid swelling and leakage (Fig. 4B, S4A, S4B). These results suggest that IAF-induced colonoid swelling is PKA-dependent.

**Figure 4.**
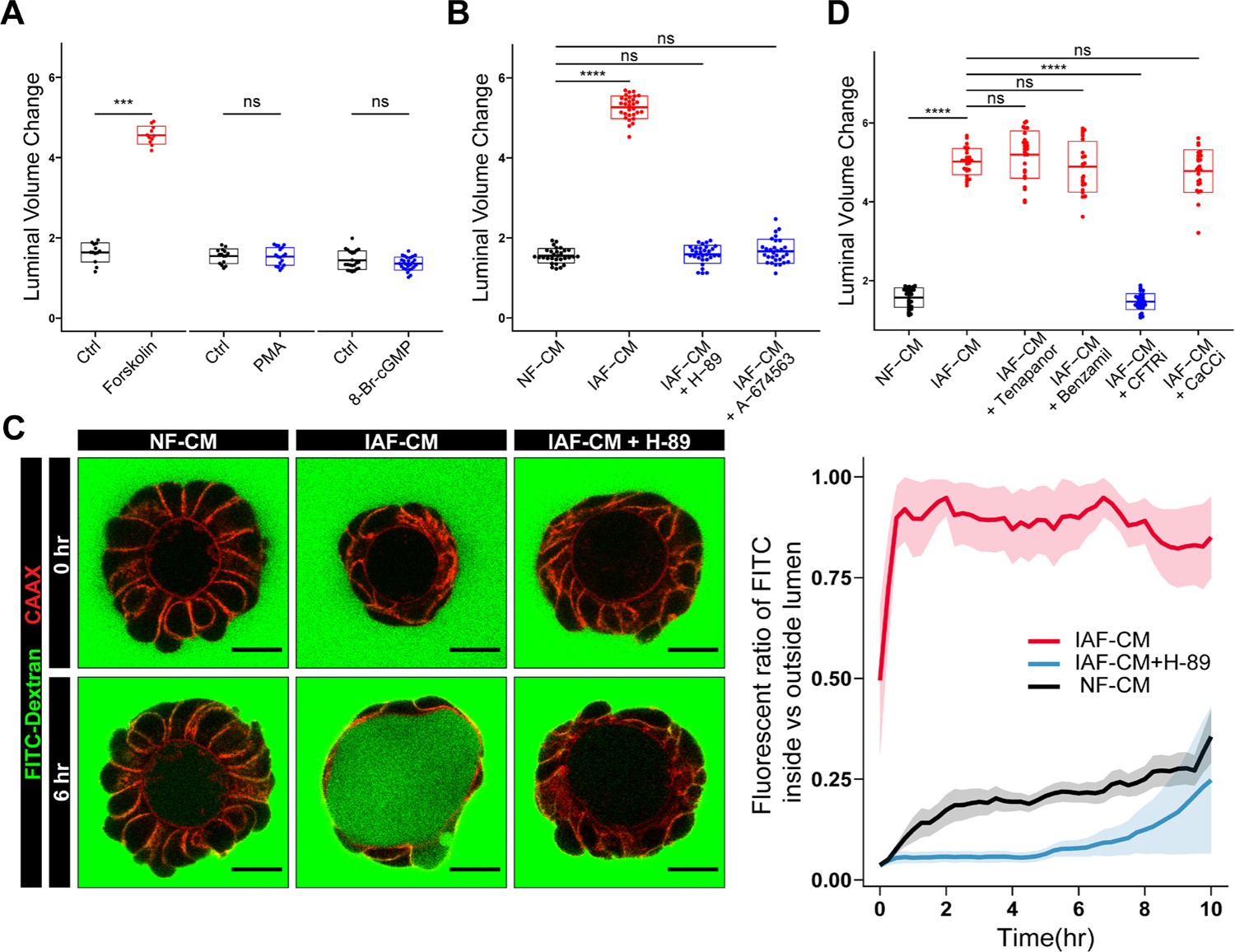
IAFs induce colonoid swelling through PKA-CFTR signaling. A. Colonoid swelling can be induced by PKA activation. Box plots show luminal volume changes of colonoids treated with PKA, PKC, and PKG agonists (Forskolin, PMA, and 8-Br-cGMP), respectively. Three independent experiments were performed with consistent results. Data shown is from a single independent experiment, with 13, 11, 15, 18, 22, and 26 colonoids measured per condition (from left to right). Each dot represents an individual colonoid; ****p < 0.0001, ns not significant, t-test, bar box represents mean ± STD. B. PKA inhibitors prevent IAF-CM-induced colonoid swelling. Box plots show luminal volume changes of colonoids cultured in IAF-CM with or without PKA antagonists H-89 and A-674563. Three independent experiments were performed with consistent results. Data shown is from a single independent experiment, with 30 colonoids measured per condition. Each dot represents an individual colonoid; ****p < 0.0001, ns not significant, t-test, bar box represents mean ± STD. C. PKA inhibitors prevent IAF-CM induced barrier leakage. Left: Representative live images of colonoids treated with NF-CM, IAF-CM, and IAF-CM with PKA inhibitor, H-89, in a FITC-Dextran permeability assay. Images shown are at the beginning of and 6-hour time point after culturing in CM. Scale bar, 20 µm. Right: Line plot showing the fluorescence ratio of FITC inside vs outside the lumen, with 6, 6, and 3 organoids quantified for NF-CM, IAF-CM, and IAF-CM + H-89, respectively. The lines show the mean and the shaded areas SEM. D. IAF-CM-induced colonoid swelling is CFTR-dependent. Box plots show luminal volume changes of colonoids cultured in IAF-CM with Tenapanor (NHE3 inhibitor), Benzamil (ENaC inhibitor), CFTR inhibitor-172 (CFTR inhibitor), or CaCC(inh)-A01 (TMEM16A family inhibitor), respectively. Three independent experiments were performed with consistent results. Data shown is from a single independent experiment, with 30 colonoids measured per condition. Each dot represents an individual colonoid; ****p < 0.0001, ns not significant, t-test, bar box represents mean ± STD.

Transepithelial fluid secretion is associated with activation and deactivation of ion channels (*43*), which may act downstream of PKA. In the colon, absorption of water is mostly regulated by sodium transporters, including sodium-hydrogen exchanger 3 (NHE3) and epithelial sodium channel (ENaC) (*40*). We treated colonoids with IAF-CM in combination with inhibitors of each (Tenapanor for NHE3 and Benzamil for ENaC). Only Benzamil reduced colonoid swelling, but not significantly (Fig. 4D, S4C, S4D). This suggests that IAF-CM induced colonoid swelling was not dependent on sodium absorption. Colon epithelial fluid secretion is known to be mostly regulated by chloride channels, namely the cystic fibrosis transmembrane conductance regulator (CFTR) and the calcium-dependent chloride channel (CaCC) (*40*). We therefore treated colonoids with inhibitors against each (CFTR(inh)-172 for CFTR and the general TMEM16 family inhibitor, CaCC(inh)-A01 for CaCC). Only the CFTR inhibitor prevented IAF-CM-induced colonoid swelling (Fig. 4D, S4D), suggesting that CFTR but not members of the calcium dependent-TMEM16 family was involved in this regulation. CFTR is a known target of PKA activation (*44*). Thus, IAF-induced organoid swelling is mediated through the PKA-CFTR axis.

### IAF-induced colonoid swelling occurs downstream of PGE2-EP4 signaling

Next, we investigated the signaling pathway upstream of PKA. Many factors can activate PKA in the context of diarrhea, including small molecules, cytokines, and neurotransmitters (*41*). To probe the molecular nature of the paracrine agent, we fractionated the conditioned media using filters of 3 kD molecular weight (MW) cutoff and treated colonoids with the top (MW > 3 kD) and bottom (MW < 3 kD) fractions (Fig. 5A). The bottom fraction from IAF-CM induced colonoid swelling more robustly than the top fraction (Fig 5A), suggesting that the major agent inducing colonoid swelling was likely to be a small molecule.

**Figure 5.**
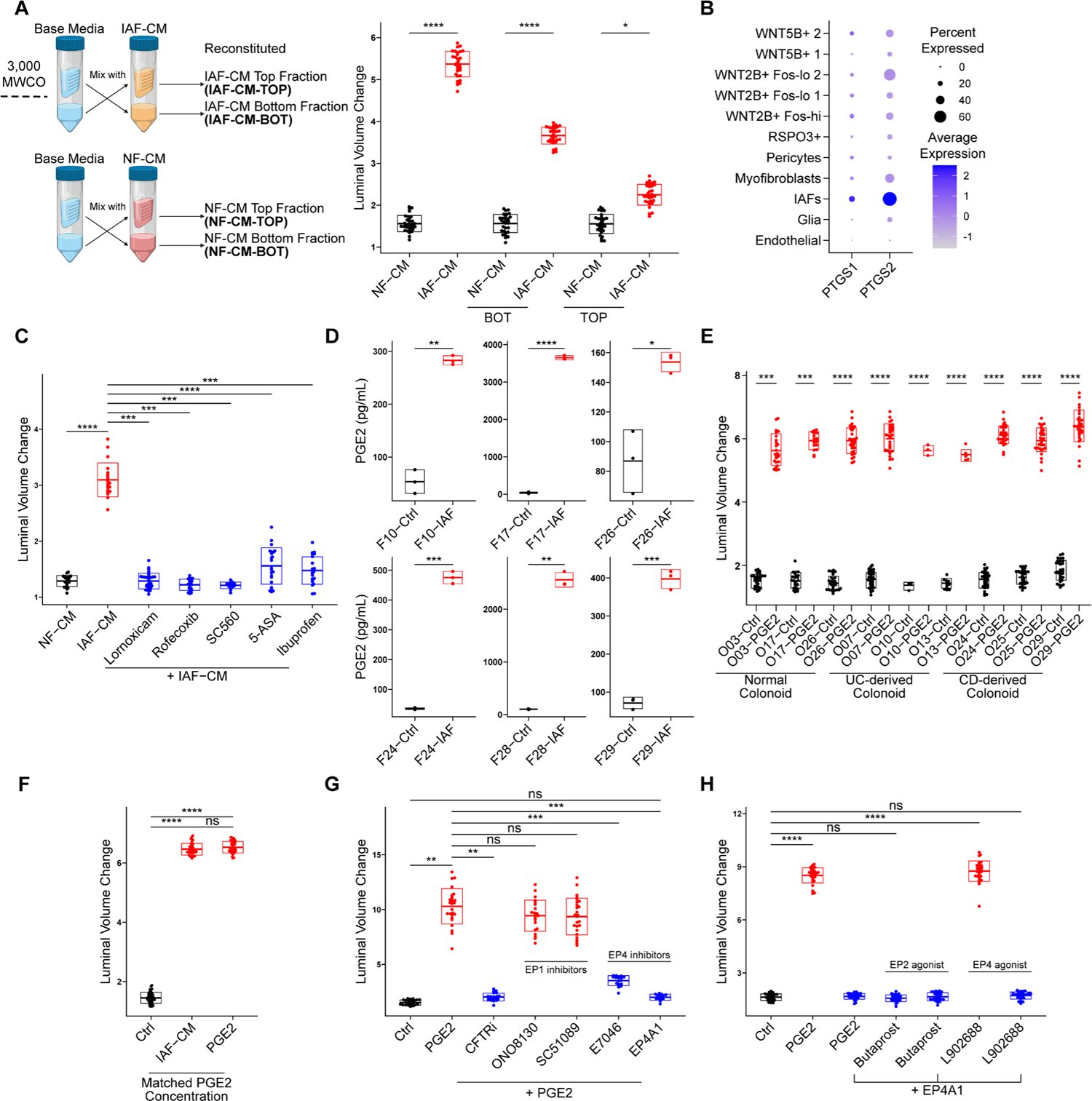
IAFs secrete PGE2 and induce colonoid swelling through EP4. A. Small molecules in IAF-CM induce colonoid swelling. Left, schematic representation of the workflow used to generate fractioned fibroblast CM for colonoid culture. 10 mL CM was fractioned into 1 mL concentrated fraction (top) and 9 mL flowthrough fraction (bot) and mixed with the complementary fraction from the base media (crossing arrows). Details are described in Methods. Right: Box plots showing the luminal volume change of colonoids cultured in top or bottom fractions of IAF-CM and NF-CM. Three independent experiments were performed with consistent results. Data shown is from a single independent experiment, with 30 colonoids measured per condition. Each dot represents an individual colonoid; ****p < 0.0001, *p < 0.05, t-test, bar box represents mean ± STD. B. *PTGS1/2* are upregulated in IAFs. Dot plot shows the expression of *PTGS1/2* (encoding COX1/2) in different subsets of patient derived fibroblasts, as described in Figure 1H. The percentage of cells expressing (dot size) and the mean expression level (dot color) of *PTGS1/2* (rows) across subsets (columns) are shown. C. IAF-CM from COX inhibitor-treated IAFs did not induce colonoid swelling. Box plot shows the luminal volume changes of colonoids cultured in IAF-CM that is derived from IAFs pre-treated with COX-inhibitors during cytokine-activation. Three independent experiments were performed with consistent results. Data shown is from a single independent experiment, with 22, 20, 30, 20, 14, 23, and 23 colonoids measured per condition (from left to right). Each dot represents an individual colonoid; ****p < 0.0001, ***p < 0.001, t test, bar box represents mean ± STD. D. PGE2 was upregulated in multiple IAF lines. Box plots show the PGE2 concentration of IAF-CM derived from six IAF lines (2 normal, 2UC, 2CD) using a PGE2 ELISA assay. Three independent experiments were performed with consistent results. Data shown is from a single independent experiment. Each dot represents a technical replicate; ****p < 0.0001, ***p < 0.001, **p < 0.01, *p < 0.05, t-test, bar box represents mean ± STD. E. PGE2 induced colonoid swelling in multiple colonoid lines. Box plot shows luminal volume changes of nine colonoids lines (3 normal, 3UC, 3CD) cultured in 10 nM PGE2. Three independent experiments were performed with consistent results. Data shown is from a single independent experiment, with 30 colonoids measured per condition except O13UC-Ctrl and PGE2 (11 and 6), O10UC-Ctrl and PGE2 (3 and 3), O03CN-PGE2 (23), O17CN-Ctrl and PGE2 (24 and 16). Each dot represents an individual colonoid; ****p < 0.0001, ***p < 0.001, t-test, bar box represents mean ± STD. F. PGE2 is the factor in IAF-CM responsible for colonoid swelling. Box plot shows luminal volume changes of colonoids cultured in IAF-CM or PGE2 at a concentration matched with the PGE2 concentration in IAF-CM (measured 2116.70 pg/mL, approximately 6.0053 nM by ELISA assay). Three independent experiments were performed with consistent results. Data shown is from a single independent experiment, with 40 colonoids measured per condition. Each dot represents an individual colonoid; ****p < 0.0001, ns not significant, t-test, bar box represents mean ± STD. G. PGE2-induced colonoid swelling is EP4-dependent. Box plots show luminal volume changes of colonoids cultured in PGE2 with different EP1 and EP4 inhibitors (EP1 inhibitors: ONO8130 and SC51089; EP4 inhibitors E7046 and EP4A1). Three independent experiments were performed with consistent results. Data shown is from a single independent experiment, with 23, 26, 21, 22, 26, 17, and 15 colonoids measured per condition (from left to right). Each dot represents an individual colonoid; ***p < 0.001, **p < 0.01, ns not significant, t test, bar box represents mean ± STD. H. EP2 is not involved in PGE2 induced colonoid swelling. Box plots show luminal volume changes of colonoids cultured in EP2 (Butaprost) or EP4 (L902688) agonists. Three independent experiments were performed with consistent results. Data shown is from a single independent experiment, with 30 colonoids measured per condition. Each dot represents an individual colonoid; ****p < 0.0001, ns not significant, t-test, bar box represents mean ± STD.

Since most small molecules activate PKA through G-protein coupled receptors (GPCRs) (*45*), we considered potential GPCR activators associated with inflammation. Among them, prostaglandins are intermediate products from the arachidonic acid (AA) pathway under the regulation of cyclooxygenase (COX) (*46*), and their production and secretion in IAFs is upregulated upon inflammation in IBD patients (Fig. S5A, S5B) (*47, 48*). To examine whether prostaglandins were involved in organoid swelling, we first confirmed increased *PTGS1/2* (encoding COX1/2) expression in IL4+IL13+TNFα activated IAFs (Fig. 5B). We then treated IAFs with COX inhibitors (including Rofecoxib, Lornoxicam, S-Ibuprofen, SC-560, and 5-Aminosalicylic Acid) during IAF induction, and collected IAF-CM for colonoid culture. Indeed, colonoids treated with COX-inhibited IAF-CM did not swell (Fig. 5C).

Prostaglandin E2 (PGE2) is the most abundant type of prostaglandin in the colon (*49*). We performed ELISA assays on IAF-CM derived from multiple fibroblast cell lines (normal and IBD-derived) and found that the level of PGE2 was significantly increased in all compared to the NF-CM controls (Fig. 5D). Treating multiple colonoid lines with PGE2 led to increased luminal volume (Fig. 5E) in a dose-dependent manner (Fig. S5C). Additionally, treating colonoids with PGE2 reduced EdU incorporation, suggesting that PGE2 contributed to the observed reduction of epithelial proliferation in IAF-colonoid co-cultures (Fig. S5D). Next, to examine whether PGE2 was sufficient to induce colonoid swelling at the concentration present in IAF-CM, we treated colonoids with a matched concentration of PGE2 as measured by ELISA in IAF-CM and found no statistical difference of luminal volume increase induced by IAF-CM or by PGE2 at the matched concentration (Fig. 5F).

Four GPCRs are known as PGE2 receptors, namely EP1, EP2, EP3, and EP4 (*50*). Among these, EP1 is a Gaq-that activates PI3K-dependent pathways (*50*). EP2 and EP4 are coupled to Gas and activate adenyl cyclase (*51*). Although both EP2 and EP4 signal through PKA, EP4 also utilizes the PI3K pathway and activates ERK1/2 (*52*). EP3 is a PKA inhibitor and not detected in colon epithelium (*53*). Because PTGER1 and 4 (corresponding to EP1 and EP4 respectively) are upregulated in inflamed epithelial cells in IBD patients (Fig. S5E), we treated colonoids with PGE2 in combination with distinct EP1 or EP4 antagonists (ONO-8130 and sc-51089 for EP1; E7046 and EP4 receptor Antagonist 1 (EP4A1) for EP4) (Fig 5G, S5F). We observed that EP4 inhibitors prevented colonoid swelling but EP1 inhibitors did not (Fig 5G, S5F). Since both EP2 and EP4 could activate cAMP (*51*), we treated colonoids with selective EP2 and EP4 agonists (Butaprost and L-902688, respectively) and observed that only the EP4 agonist induces organoid swelling (Fig 5H, S5G). These data show that IAF-induced colonoid swelling is mediated by PGE2 activation of EP4.

Lastly, to determine whether PGE2 signaling was sufficient to cause epithelial barrier disruption, we treated colonoids with 10 nM PGE2, a concentration at the higher end among different batches of IAF-CM (Fig. S5H). The PGE2-treated colonoids showed increased luminal FITC-dextran leakage, which was rescued by an EP4 inhibitor (Fig. S5H). Treating colonoids with IAF-CM showed slightly higher permeability than matched PGE2 (Fig. S5I, S5J). These data suggest that PGE2 is critical for the IAF-induced barrier disruption phenotype, but additional factors may also be involved (*54*).

### IAF-induced colonoid swelling increases DNA damage and mitotic errors which are mitigated by EP4 inhibition

Since rapid changes in osmotic or mechanical stress can increase DNA damage and compromise mitotic fidelity (*55-59*), we hypothesized that IAF-induced epithelial swelling, which dramatically alters cell and nuclear shape, could increase mitotic errors and DNA damage, which are causes of CIN associated with CAC initiation and progression (*60*). To test this, we first monitored cell divisions by live-cell imaging using normal colonoids expressing both H2B-mNeonGreen and mCherry-CAAX. Colonoids treated with IAF-CM showed increased mitotic errors, including chromosome bridges and lagging chromosomes (Fig. 6A, 6C). To confirm that this phenotype can be caused by PGE2 signaling, we treated colonoids with 10 nM PGE2 and observed a similar increase in mitotic errors (Fig. 6B). Time-lapse imaging of PGE2-treated colonoids showed many examples of grossly misaligned chromosomes during mitosis of swollen and distorted epithelial cells (Fig 6D, S6A, supplemental movie 4). Additionally, 53BP-1, a marker of the DNA damage response, accumulated and persisted at sites where chromatin bridges were severed during cytokinesis (Fig. 6E, supplemental movie 5). Immunofluorescent staining of colonoids with γH2AX, a marker of double-stranded DNA breaks, confirmed increased DNA damage in both IAF-CM and PGE2 treated colonoids compared to controls (Fig. 6F, 6G).

**Figure 6.**
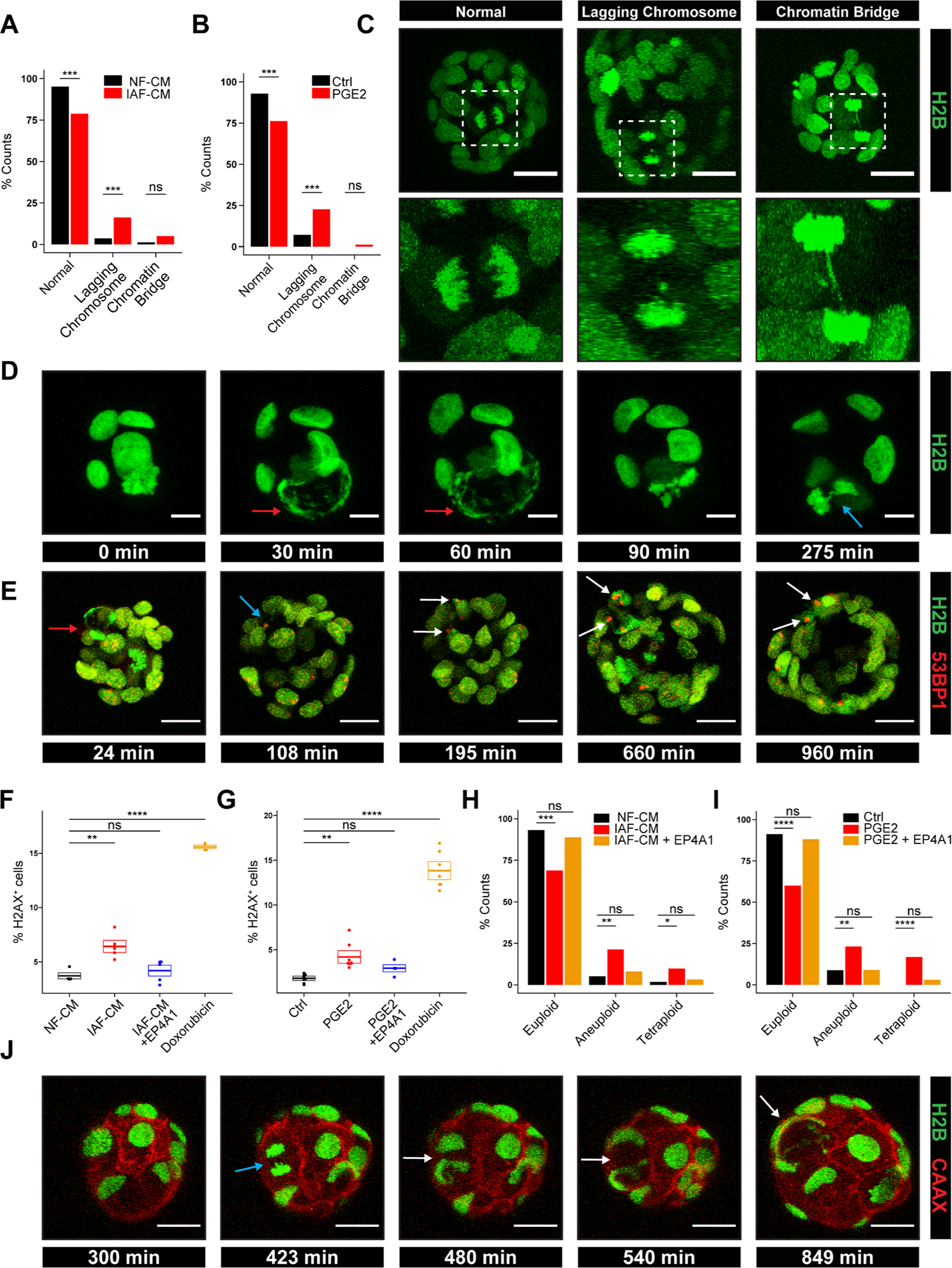
PGE2 negatively affects mitotic fidelity and increases the incidence of DNA damage. A. Colonoids exhibit increased mitotic errors when cultured in IAF-CM. Bar plot shows the fraction of colonoid cells that undergo chromatin bridge, lagging chromosomes, or normal mitosis when cultured in IAF-CM or NF-CM. n = 82 and 86 for IAF-CM treated and NF-CM treated conditions, respectively. ***p < 0.001, ns not significant, Fisher’s exact test. B. Colonoids exhibit increased mitotic errors when cultured in PGE2. Bar plots showing the fraction of colonoid cells that undergo chromatin bridge, lagging chromosomes, or normal mitosis when cultured with PGE2 or base media (control). n = 84 and 67 for PGE2 treated and control conditions, respectively. Fisher’s exact test. C. Colonoids undergoing normal or erroneous mitosis. Representative images of colonoid cells undergoing mitosis with chromatin bridge, lagging chromosome, or normal mitosis. Scale bar, 20 µm. D. Swelling of individual colonoid cells affects mitosis. Representative live cell images showing a cell in a colonoid in M phase at the beginning of PGE2 treatment. PGE2-induced cell swelling disrupted alignment of chromosomes (red arrows) and led to erroneous mitosis (blue arrow). Scale bar, 10 µm. E. Mitotic error is a cause of DNA damage in colonoids. Representative live cell images showing that mitotic errors led to DNA damage. The red arrow shows a PGE2-treated dividing cell with a chromatin bridge that led to DNA damage (blue arrow). 53BP1 foci persisted for more than 12 hrs (white arrows). Scale bar, 20 µm. F. IAF-CM increases the incidence of DNA damage in colonoids. Dot plot shows the percentage of H2A.X positive cells after 6 hrs of IAF-CM. 0.5 nM Doxorubicin was used as a positive control. n = 4, 5, 5, 4 for NF-CM, IAF-CM, IAF-CM+EP4A1, and doxorubicin, respectively. ****p < 0.0001, **p < 0.01, t-test, bar box represents mean ± STD. G. PGE2 increases the incidence of DNA damage in colonoids. Dot plot shows the percentage of H2A.X positive cells after 6 hrs of 10 nM PGE2 treatment. 0.5 nM Doxorubicin was used as a positive control. n = 6, 5, 3, 6 for control, PGE2, PGE2+EP4A1, and doxorubicin, respectively. ***p < 0.001, **p < 0.01, t-test, bar box represents mean ± STD. H. IAF-CM-treated colonoids have more aneuploidy and tetraploidy. Box plot shows the percentage of chromosome counts of colonoids treated with NF-CM, IAF-CM, or IAF-CM + EP4A1 in the metaphase chromosome spread assay. 58, 61, and 62 spreads were counted for NF-CM, IAF-CM, and IAF-CM+EP4A1 conditions, respectively. ***p < 0.001, **p < 0.01, *p < 0.05, ns not significant, Fisher’s exact test. I. PGE2-treated colonoids have more aneuploidy and tetraploidy. Box plot shows the percentage of chromosome counts of colonoids treated with base media (control), PGE2, or PGE2+EP4A1 in the metaphase chromosome spread assay. 57, 125, and 67 spreads were counted for control, PGE2, and PGE2+EP4A1 conditions, respectively. ****p < 0.0001, ***p < 0.001, ns not significant, Fisher’s exact test. J. PGE2 induces tetraploidy via cytokinesis failure. Representative live cell images of a colonoid cell that underwent normal metaphase (blue arrow) but failed cytokinesis (white arrows). Scale bar, 20 µm.

To evaluate whether mitotic errors led to aneuploidy, we performed chromosome counting using metaphase spreads (Fig. S6B). We observed a significant increase in aneuploid and tetraploid cells in colonoids treated with IAF-CM (Fig. 6H, S6C). Similar results were observed when treating colonoids with PGE2 (Fig 6I, S6D). Live cell imaging revealed that tetraploidy can arise from failed cytokinesis, which could be disrupted due to cell swelling (Fig. 6J). Interestingly, we observed the formation of large intracytoplasmic vacuoles (>5 µm) that displaced cytoplasmic contents (Fig. S6E, S6F, supplemental move 3-5) in swollen epithelial cells, which could also interfere with cell division. Importantly, treating colonoids with an EP4 inhibitor prevented both increased DNA damage and the induction of aneuploidy and tetraploidy caused by IAF-CM or PGE2 (Fig. 6F-6I). Thus, IAF-induced colonoid swelling is a source of genome instability that can be mitigated by EP4 inhibition.

## Discussion

Although numerous studies have implicated IAFs in IBD pathogenesis, none have directly queried the interaction between IAFs and colon epithelium (*15, 22, 61, 62*). Here, we develop an *in vitro* method of co-culturing human colon derived IAF and colon epithelium in order to investigate the role of IAFs in instigating and perpetuating epithelial dysfunction relevant to IBD biology. Using scRNA sequencing, live cell imaging, and biochemical assays, we found that IL4+IL13+TNFα-induced fibroblasts recapitulate key features of IAFs *in vivo*. These cytokine-activated IAFs cause colonoid swelling and barrier disruption in a PKA and CFTR-dependent manner downstream of PGE2-EP4 signaling. IAF-induced epithelial swelling leads to DNA damage and mitotic errors, which can be mitigated by inhibiting the PGE2 receptor EP4. Our findings thus shed light on the cellular and molecular mechanism of IAFs and the importance of prostaglandin signaling in IBD pathogenesis.

Although upregulation of PGE2 in IBD tissue has been recognized for decades (*63, 64*), its major cellular source and role have been unclear and even controversial. Previous studies have reported macrophages, monocytes, mesenchymal stem cells, and T cells as putative sources of PGE2 (*65-69*). However, none of those cell types alone produce the same level of PGE2 measured in IBD patients (*70, 71*). Our data show that IAFs can produce nanomolar concentrations of PGE2, suggesting that they may be a major source of PGE2 in IBD. Regarding the function of PGE2, previous studies in IBD suggest that PGE2 can be both pro-inflammatory and anti-inflammatory in the context of different recipient cells (*66-69, 72, 73*). Our data suggest that PGE2 has a pro-inflammatory role in the context of epithelial cells, leading to epithelial swelling, crypt distortion, disrupted barrier function, mitotic errors, and DNA damage. Interestingly, in addition to IBD, IAFs and PGE2 are also associated with other chronic inflammatory diseases such as idiopathic pulmonary fibrosis (IPF) (*74*), rheumatoid arthritis (RA) (*75*), and chronic kidney disease (CKD) (*76*). The pathway that we have identified may help explain the role of IAFs in the pathogenesis of these diseases.

PGE2 is known to promote isosmotic fluid secretion by stimulating chloride secretion while blocking sodium absorption (*77*). This action is accompanied by water secretion across the epithelium. Our results show that tight junctions were overall intact during IAF-induced colonoid swelling, suggesting that fluid flow was trans-cellular rather than intercellular. This is consistent with our observation that individual cells in colonoids could swell when exposed to IAF-CM. Cell swelling is likely to be a consequence of unbalanced or uncoordinated fluid influx vs efflux. While most cells in colonoids did not undergo dramatic swelling, even one or few cells in a colonoid undergoing dramatic swelling and rupture could lead to local disruption of epithelial barrier, as shown by live imaging. In mitotic cells that underwent swelling, the spindle and chromosome alignment appeared grossly abnormal, leading to segregation defects and DNA damage. This may be consistent with previous reports that cell compression and shape distortion disrupt spindle morphogenesis and lead to mitotic errors (*55, 56*). Additionally, DNA damage can occur when the nucleus is compressed or distorted (*57, 59*). Indeed, we observed many cases in live-cell imaging assays where cell nuclei are flattened or deformed into lobular shapes (Fig. S6F).

Our study provides insights into the association of persistent IAFs and treatment-refractory IBD. We show that multiple cytokine cocktails, even without TNFα, can lead to varying IL13RA2 expression and IAF phenotypes (Fig. 1D). Thus, resilience of the IAF phenotype in the setting of variable cytokine milieus and pharmacologic TNFα blockade could contribute to heterogeneity of patient disease trajectories and treatment responses. For example, IL1β+TNFα-activated IAFs showed stronger chemotaxis-associated signatures (Fig. 1K, 1L), which could promote more robust immune infiltration into the lamina propria. These data highlight the potential value of combinatorial immunomodulatory therapeutic approaches in IBD.

Another potentially translatable finding of our study is that EP4 is a critical mediator of the IAF-epithelial interaction. Mutations in the EP4 encoding gene *PTGER4* are associated with both UC and CD in a GWAS study (*78*). We found that blocking EP4 prevents IAF-induced epithelial swelling, barrier disruption, DNA damage, and mitotic errors. Others have suggested a role for EP4 antagonists in modulating regulatory T cells and gut microbiome in chronic intestinal inflammation (*76*). Our data identify another key role for PGE2-EP4 antagonism in promoting epithelial homeostasis. As such, EP4-antagonists may be useful therapeutic agents to treat both acute and chronic sequelae of IBD. To date, EP4 antagonists have not been used in IBD-associated clinical trials but are under active clinical trials for treating solid tumors in combination with PD-1 blockades. Several of these trials have completed dose escalation and have entered phase 1b (NCT04443088) or phase 2 (NCT02538432, NCT03696212), suggesting that these agents may be exploited for promoting epithelial barrier integrity and mitigating CAC risk.

Although our study provides a mechanistic understanding of how human IAFs affect colon epithelium *in vitro* using patient-derived fibroblasts and colonoids, our study was not extended to observing their interactions directly *in vivo*. However, in a previous report, knocking out *Ptgs2* (encoding *COX2*) in fibroblasts reduced the level of PGE2 in mice and prevented tumor initiation in the azoxymethane (AOM) colon cancer model (*72*). Our results provide a possible explanation for this observation in a patient-derived culture system. We also note that EP4 was reported to suppress TNFα-induced intestinal epithelial necroptosis by inhibiting MLKL oligomerization and membrane translocation (*79*), suggesting that PGE2 signaling has multiple roles in IBD.

## Materials and Methods

### Patient sample collection

Fresh colonic tissue was harvested from patient specimens (CD, UC, and unaffected normal) within thirty minutes of surgical resection, with informed consent under approved Johns Hopkins School of Medicine Institutional Review Board protocol IRB00125865. All specimens were carefully evaluated, annotated, and harvested by an expert gastrointestinal pathologist (T.L.). Clinical information and metadata are summarized in Supplemental Table 1. A corresponding formalin-fixed paraffin embedded (FFPE) sample was taken for each fresh sample for histological evaluation (Supplemental Table 1). Fresh samples were immediately placed into complete DMEM (cDMEM, 10% HI-FBS + 1x antibiotic-antimycotic in DMEM high glucose) and transported at 4°C.

### Colonoid and fibroblast line generation

Samples were transported to the lab within 1 hour of surgical resection. They were briefly washed in ice cold PBS containing 1x Pen/Strep and trimmed into smaller pieces (< 1mm^2^). Samples were washed extensively using ice cold HBSS until the supernatant turned clear. For post-processing, most surgical samples were immediately stored in 90% heat inactivated FBS + 10% DMSO and cryopreserved in liquid nitrogen. Remaining samples were used to generate colonoids and/or fibroblasts.

For colonoid generation, samples were transferred into a 50 mL conical tube containing 10 mL stripping buffer (5mM EDTA + 1x Pen/Step + 2% FBS + 1x HEPES in HBSS without Ca^2+^/Mg^2+^). Sample pieces were shaken on an orbital shaker at 37°C for 20 min at 200 rpm to release crypts. Released crypts were collected to generate colonoid lines as previously described (*34*).

For fibroblast isolation, minced tissue was transferred into a 50 mL conical tube containing 10 mL digestion buffer (0.05 mg/mL Liberase TH + 1x Pen/Step + 2% FBS + 1x HEPES in HBSS with Ca^2+^/Mg^2+^). Sample pieces were shaken on an orbital shaker at 37°C for 60 min at 200 rpm. Then, digested cells were filtered through a 40 µm strainer and centrifuged at 200 xg for 5min at 4°C. After removing the supernatant, red blood cells were removed by adding 2 mL ACK lysis buffer for 2 min at R.T. followed by another round of centrifugation. Lastly, cells were resuspended in cDMEM and seeded in 6-well plates at 2M cells/mL. 48 hrs post seeding, the media was changed to remove both dead and suspension cells. The media was then changed once every three days and cells were passaged when they reached 90% confluency. All fibroblasts were used for experiments within the first 10 passages.

### Single Cell RNA sequencing of patient-derived fibroblast lines

For fibroblast scRNA-seq experiments, the Chromium Next GEM single cell 3’ HT reagent kit V3.1 was used for sample preparation. After QC, sequencing was performed using a S4 flow cell in a NovaSeq 6000 system. The targeted read depth was 50 K reads per cell and 2 K reads per CMO barcode. After sequencing, data were uploaded to the 10x cloud analysis server and analyzed using Cell Ranger 7.0.1. Further analysis was performed in R (4.3.0) using Seurat (4.2.0), monocle3 (1.3.1), followed by the standard data processing and analysis workflow.

Reference mapping was performed using a slightly modified protocol from the original publication (*29*). In brief, Seurat objects were normalized using SCTransform and anchors were calculated using the standard PCA transformation (UCatlas as reference and our fibroblast cell dataset as the query dataset) to find and transfer matched cell type labels. Then, the query dataset was projected onto the UMAP structure. Wilcoxon rank-sum test was used to find markers for each subset. To compare the DEGs between IL1β+TNFα IAFs vs IL4+IL13+TNFα IAFs, pseudobulk analysis was performed in Libra (*80*), using edgeR (3.17) LRT method. Gene set enrichment analysis (GSEA) was performed using rankings and annotated using org.Hs.eg.db (3.17.0).

### Colonoid related culture methods

#### General colonoid culture

Colonoids were cultured and passaged in organoid expansion media (including EGF, Wnt3A, R-spondin1, and Noggin, aka EWRN media). For the first three days after seeding, 10 µM Y-27632 was added to avoid anoikis. Then, expansion media was changed once every two days and colonoids were passaged weekly. For all experiments, colonoids were used before passage 25. Experiments with expanding colonoids were performed on day 4 after single-cell seeding. All small molecules drugs were reconstituted in DMSO and stored at −20 °C. Unless otherwise specified, the concentration of the chemicals was as follows: Forskolin (10 µM), H-89 (50 µM), A-674563 (10 µM), Tenapanor (10 µM), Benzamil (10 µM), CFTRi (10 µM), CaCCi (10 µM), PGE2 (10 nM), ONO8130 (10 µM), SC51089 (10 µM), E7046 (50 µM), EP4A1 (10 µM), Butaprost (10 nM), and L902688 (10 nM).

#### Colonoid differentiation

For experiments using differentiated colonoids, colonoids were switched to differentiation media (expansion media without Wnt3A and SB202190, a p38 inhibitor) on day 4 after single-cell seeding and maintained for 4 additional days. Media was renewed once every two days. Colonoid differentiation was visually confirmed by observation of budding phenotypes, as described previously (*34*).

#### Transwell colonoid-fibroblast co-culture

Colonoids were mixed in 20 µL Matrigel, seeded in the Transwell (0.4 µm PET membrane, Corning Cat#353095), and settled in a glass-bottom 24-well plate (Cellvis, Cat# P24-1.5H-N). 100 µL and 600 µL of expansion media were added into the transwell and the plate, respectively. Colonoids were cultured for 4 days before dissociated fibroblasts were added. Imaging was then performed using the Nikon TiE microscope, as described in the imaging section.

#### Colonoid culture for transmission electron microscopy

Colonoids were mixed in 10 µL Matrigel, loaded in a sterilized specimen carrier A (specimen carrier, 6 mm, 0.1/0.2 mm) (Technotrade 1190-100), and cultured in a 24-well plate for 4 days. Expansion media was then switched to fibroblast-conditioned media for 6 hours followed by high pressure freezing, as described in HPF section.

### Fibroblast related culture methods

#### Cytokine induction

Fibroblasts were seeded at about 20% confluency. Two days after seeding, fibroblasts were treated with cytokines for 4 days. Then, cells were collected for downstream assays, including western blot, co-culture, and conditioned media generation. For cytokine induction, unless otherwise stated, all cytokines were reconstituted as 1,000x working stocks in cDMEM at 10 µg/mL except TNFα (50 µg/mL) and stored at −20°C prior to use.

#### Generation of fibroblast conditioned media (CM)

After cytokine induction by IL4+IL13+TNFα, fibroblasts were washed three times in PBS to remove remaining cytokines and supplied with organoid culture media (expansion or differentiation media) for 1 day. Conditioned media was then collected and centrifuged at 600 xg for 5 min to remove cell debris. CM was stored at −20 °C for long term storage and at 4 °C for immediate use within a week.

#### Generation of COX inhibitor-treated IAF-CM

During cytokine induction, COX inhibitors were added together with IL4+IL13+TNFα. Then, drugs were washed away, and conditioned media was generated using the same approach as described above. COX inhibitors were reconstituted in DMSO and stored as 1000x working stocks at −20°C prior to use. The default working concentrations of each were: Lornoxicam (50 µM), Rofecoxib (50 µM), S-Ibuprofen (50 µM), SC-560 (50 µM), and 5-ASA (200 µM).

### Ultrafiltration

Ultrafiltration was performed using a 3,000 molecular weight cutoff (MWCO) PES concentrator (Pall Lab, Cat#MAP003C36). 10 mL of media was loaded to the concentrator and centrifuged at 5000 xg for 2 hours at 4°C to generate an approximately 1 mL concentrated fraction (top fraction) and a 9 mL flowthrough fraction (bottom fraction). After ultrafiltration, the concentrated fraction (top fraction) of the conditioned media was reconstituted with the flowthrough fraction (bottom fraction) of the base media; the flowthrough fraction (top fraction) of the conditioned media was reconstituted with the concentrated fraction (bottom fraction) of the base media.

### Confocal imaging

Confocal Imaging was performed using a Zeiss LSM880-Airyscan FAST microscopy equipped with an incubation chamber. Colonoids were seeded on 8-well chambered coverglass (Nunc LabTek II, Cat# 155360 or ibidi Cat# 80827) and imaged 4 days after single-cell seeding. For live-cell imaging, cells were incubated at 37°C with 5% CO2 and imaged using a 40x/1.20 C-Apo water objective with Zeiss Immersion Oil W 2010 media. For cell division assays, colonoids were imaged once every 3 min with 30-50 1.0-1.5 µm z-slices. For FITC-dextran permeability assays, colonoids were imaged once every 15 min with 30-40 1.5-2.0 µm z-slices. For fixed cell imaging, the airyscan mode was used with default settings (zoom factor = 2.0, 40x water objective for proliferating colonoids and 20x air objective for differentiated colonoids). Images were analyzed using ImageJ (version 1.53C).

### Imaging and quantification of colonoid swelling

The colonoid swelling assay was performed using a Nikon TiE microscope equipped with an incubation chamber (37°C, 5% CO2). Colonoids were cultured in 24-well glass bottom black plates (Cellvis, Cat# P24-1.5H-N) for 4 days and imaged using a 10x objective. Once the areas of interest were selected (3 areas per well, each with dimensions of = 1311.20 µm x 1328.60 µm), culture media was changed to designated media and imaged every 15 min for 12-48 hrs. Images were analyzed using ImageJ. The level of swelling was quantified by comparing the fold change of the luminal area before and after the treatment (at 24 hours post treatment, unless otherwise described). Quantified data were organized and plotted using RStudio (2022.12.0 Build 353).

### Lentivirus production and generation of modified colonoid lines

Lentiviruses were generated using pMD2.G, psPAX2, and corresponding lentiviral transfer constructs. We followed the forward transfection protocol of the lipofectamine 3000 reagent (Invitrogen, Cat# L3000008) with some minor changes. In brief, HEK293FT cells were passaged at 1:4 ratio 36hrs before transfection. Cells were co-transfected with the lentiviral transfer plasmid, packaging plasmid (psPAX2), and envelop plasmid (pMD2.G) at a 1:1:1 molar ratio using Lipofectamine 3000 for 6 hrs. Then, media containing lentivirus was collected and concentrated 100-fold using the Lenti-X Concentrator (Takara Bio, Cat# 631231). Lentiviral titers were determined using the qPCR Lentivirus Titration Kit (abm, Cat# LV900). Lentiviruses were aliquoted into cryogenic tubes (100 µL per tube per transfection), snap frozen in liquid nitrogen, and stored at −80 °C.

To generate modified colonoid lines, colonoids were dissociated into single cells and resuspended in 1 mL colonoid expansion media containing 100 µL concentrated lentivirus, 10 µM Y-27632 and 0.8 µg/mL polybrene. Single cells were spinfected at R.T. for 1 hr at 70 xg. Cells were then incubated for 6 hrs (37°C, 5% CO2) with gentle agitation once per hour. Lastly, cells were rinsed with base media and embedded in Matrigel (1,000 cells/µL, 20 µL per well in a 24 well plate). Modified colonoids were cultured normally for the first passage. At the beginning of the second passage, puromycin (1 µg/ml) and/or blasticidin (5 µg/ml) were added to media depending on the selection gene(s). For colonoids with some fluorescent reporters, dissociated single-cell colonoids were bulk sorted using a Sony SH800 cell sorter at the end of the third passage to further increase population purity.

### EdU incorporation

Colonoids (4 days after single-cell seeding) were treated with 10 µM EdU for 6 hours before dissociation and staining using the Click-It EdU Alexa Fluor 488 Flow Cytometry Assay kit (ThermoFisher, Cat# C10425).

### Immunofluorescence staining of colonoids (*in situ*)

Colonoids were washed three times in PBS and fixed in 4% PFA for 10 min at 37 °C. After fixation, colonoids were washed 3 times using PBS for 10 min each at RT. Then, colonoids were permeabilized using 0.5% Triton X-100 in PBS for 1 hour at RT and blocked in blocking buffer (10% BSA + 0.1% Triton X-100 in PBS) for 1 hour at RT. Colonoids were stained with primary antibody in staining buffer (1% BSA + 0.1% Triton X-100 in PBS) at 4°C overnight. After primary antibody staining, colonoids were washed 3 times in PBS and stained with secondary antibody for 4 hours at RT in staining buffer. Finally, colonoids were washed and left in PBS at 4°C until imaging.

### Western Blot

Protein lysis buffer was made fresh before each experiment and kept on ice. Lysis buffer included 142.8ul 7x protease/phosphatase inhibitor stock (mixture of Roche Cat#11836153001 and Cat#4906845001), 100ul 10x RIPA buffer, 50ul glycerol, 10ul 10% SDS, and 697.2ul ddH2O. Colonoids (25 µL per well in 24-well plate) and fibroblasts (150 µL per well in 6-well plate) were collected in 1.5 mL Eppendorf tubes and lysed in protein lysis buffer for 30 min with brief vortexing once every 10 min. Lysed cells were further dissociated by sonication at 70 V for 5 min at 4°C and then centrifuged for 10 min at 4°C at 17,000 xg and supernatant was collected. Protein concentrations were measured using a BCA analysis kit (ThermoFisher Cat#23225). After normalizing protein concentration, proteins were mixed with 4x loading buffer and 50 mM DTT. For Western blots, NuPAGE 4-12% Bis-Tris gels (10/12/15 wells) and MES-SDS (ThermoFisher Cat# B000202) running buffer were used with default settings. For protein transfer, the iBlot 2 gel transfer system (ThermoFisher Cat# IB21001) was used with default settings. For imaging, ECL substrate (BioRad, Cat# 170-5060) was used as the detection reagent and the Odyssey XF imaging system (LI-COR Biosciences) was used to record images. Restore Plus Western blot stripping buffer (ThermoFisher Cat# 46430) was used for membrane stripping.

### mRNA extraction and qPCR

Trizol (ThermoFisher Cat#15596026) was used to collect mRNA. mRNA was extracted using the RNA Mini Kit (ThermoFisher Cat#12183018A) with column DNase treatment (Qiagen Cat#79254). After elution, mRNA concentrations were measured using a NanoDrop and normalized to approximately 125 ng/µL. Reverse transcription was performed using SSIV VILO mater mix (ThermoFisher Cat#11756050). The cDNA was diluted 20-fold in ddH2O and mixed with SYBR Green master mix (QuantaBio Cat#95073-05K) and designated qPCR primers. Each reaction was 10 µL and each test was performed in triplicate. ACTB was used as the control gene and untreated fibroblasts were used as control samples. The delta-delta CT method was used for data analysis and Log-2 fold change (Log2FC) was used for data presentation.

### Karyotyping

Colonoids were treated with expansion media containing 10% Colcemid (1 µg/ml) for 8 hours and dissociated into single cells using TrypLE Express. Single cells were resuspended in 6 ml hypotonic solution (0.56% pre-warmed KCl) and incubated in a 37 °C water bath for 10 min. Then, cells were pre-fixed by adding 1.5ml fixation buffer (Methanol:Glacial acetic acid = 3:1) and incubating in water bath for 5 min. Then, cells were centrifuged at 300 xg at R.T. for 5min. After removing the supernatant, cells were fixed in 6 mL fixation buffer and incubated in a water bath for 10 min. After centrifugation, cells were resuspended in 6 mL fixation buffer and sent for centrifugation immediately without incubation. Resuspended cells were dropped onto a heat-moisturized imaging slide (2 drops per slide) and incubated on a heater for 30 min at 75°C. Lastly, 20 µL of mounting media with DAPI was loaded evenly onto the slide and a #1.5 22×50mm cover glass was used to cover the spreads. Spreads were counted using a Nikon TiE microscope 63x oil objective and 1.5x focal reducer. At least 50 counts were recorded per condition.

### Flow cytometry

Cells were filtered and analyzed using the Attune NxT flow cytometer (Thermo Fisher Scientific). Results were further processed using FlowJo (BD, Version 10) for raw data and flow plots. Statistical analysis was then performed with R (version 4.2.2).

### High pressure freezing and freeze substitution

Colonoids were cultured in specimen carrier A (specimen carrier, 6 mm, 0.1/0.2 mm) (Technotrade 1190-100) and frozen using a high pressure freezer (EM ICE, Leica Microsystems). 20% BSA dissolved in colonoid culture medium was used as cryo-protectant. Specimen carrier A with the cells facing up was mounted in the sample holder and enough cryoprotectant was added to cover the cells. Another specimen carrier with a flat side (specimen carrier, 6 mm, 0.3 mm/flat) (Technotrade 1191-100) was placed on top of the specimen carrier A and a 200 µm spacer ring (Leica) was placed on top. The entire assembly was placed in between the half cylinders and frozen using the high-pressure freezer. Frozen samples were dropped in a liquid nitrogen storage container and transferred to an automated freeze substitution (AFS2. Leica Microsystems) unit, keeping the samples under liquid nitrogen. Freeze substitution was performed using two fixatives, fixative I containing 1% glutaraldehyde (Electron Microscopy Sciences, 16530), 0.1% tannic acid (Sigma, 403040-100G), and fixative II containing 2% osmium tetroxide (Electron Microscopy Sciences, 19132) both prepared in anhydrous acetone. Right after freezing samples were placed in AFS2, in a universal sample container (Leica Microsystems) containing fixative I, pre-chilled at −90 °C and left at −90 °C for 40h. After that samples were washed with pre-chilled acetone (−90 °C) for 5 times, 30 minutes per wash. After the last acetone wash freeze substitution solution II, prechilled inside AFS2 was added to the samples. The following steps were performed to complete the freeze substitution process: −90 °C for 41 h, −90 °C to −20 °C in 14 h, −20 °C for 12 h and −20 °C to 4 °C in 2h, and samples were held at 4 °C until further processing. Sample containers were covered with a clear film to prevent evaporation.

### Sample preparation for electron microscopy

Following freeze substitution, fixatives were washed with anhydrous acetone 5 times, each wash for 20 minutes. 100% Epon Araldite was prepared (Epon 6.2 g, Araldite 4.4 g, DDSA 12.2 g, BDMA 0.8 ml) (Epon-Araldite kit, Ted Pella, 18028) and 30%, 70% and 90% dilutions were prepared from 100% Epon with acetone. Samples were infiltrated with 30% and 70% Epon solutions for at least 2 h and 90% overnight. The following day, samples were transferred to freshly prepared 100% Epon and Epon solution was changed two times. At the final step the carriers containing the colonoids were placed at the bottom of a BEEM capsule (Electron Microscopy Sciences, 102096-558) such that the cells were facing up and the capsule was filled with 100% Epon. The samples embedded in Epon were cured at 60 °C for 48 hr. After the resin was cured 70-90 nm sections were cut using an ultramicrotome (EM UCT, Leica Microsystems) and collected on a 2×1 mm copper slot formvar coated grid.

### Transmission electron microscopy and image analysis

Samples were imaged on a Hitachi 7600 TEM equipped with an AMT XR80 camera with an AMT capture V6 at 80 kV at typically 30,000x magnification. Intercellular junctions were imaged and analyzed using ImageJ.

### Quantification and statistical analysis

Please refer to Figure Legends or the corresponding Methods for the description of sample size and statistical details. Statistical analysis was performed using R-Studio (2022.12.0 Build 353, R version 4.2.2). For all statistical tests, p < 0.05 was used as the cutoff to indicate significance.

## Acknowledgments

We thank Linda D. Orzolek and Tyler J Creamer from the Johns Hopkins Single cell & Transcriptomics core for conducting single-cell RNA-sequencing. We thank Hao Zhang from Johns Hopkins Bloomberg Flow Cytometry and Immunology Core for performing cell sorting. We thank LaToya Roker and Hoku West-Foyle from the Johns Hopkins Microscope Facility for suggestions with electron and confocal microscopy. We thank Bertram Bleck from Takeda Pharmaceuticals for insightful discussions. We thank Haiyang Wang from the Mechanical Biology Institute of the National University of Singapore for insightful discussions. We thank Sreekumar Ramachandran and Josh McNamara from the Johns Hopkins department of cell biology for insightful discussions.

## Author contributions

Conceptualization, Y.D. and R.L.; Methodology, Y.D., T.C.L., M.D.;

Investigation, Y.D., B.A.J., L.R., M.Z., A.Z.L., S.R., I.C., J.Z., B.S., N.Z.; Resources, T.C.L., S.W., P.S.;

Manuscript Writing, Y.D., T.C.L., R.L.;

Manuscript Review and Editing: Y.D., J.Z., T.C.L., M.D., R.L.; Visualization, Y.D.;

Supervision, T.C.L., R.L.;

Funding Acquisition, T.C.L., and R.L.

## Data and materials availability

The scRNAseq data is deposited to the NCBI GEO repository (GSE237034). All data are available in the main text or the supplementary materials.

